# Leveraging large language models for data analysis automation

**DOI:** 10.1101/2023.12.11.571140

**Authors:** Jacqueline A Jansen, Artür Manukyan, Nour Al Khoury, Altuna Akalin

## Abstract

Data analysis is constrained by a shortage of skilled experts, particularly in biology, where detailed data interpretation is vital for understanding complex biological processes and developing new treatments and diagnostics. To address this, we developed `mergen`, an R package that leverages Large Language Models (LLMs) for data analysis code generation and execution. Our primary goal is to enable humans to conduct data analysis by simply describing their objectives and the desired analyses for specific datasets through clear text. Our approach improves code generation via specialized prompt engineering and error feedback mechanisms. In addition, our system can execute the data analysis workflows prescribed by the LLM providing the results of the data analysis workflow for human review. We evaluated the performance of this data analysis system using various data analysis tasks. Our evaluation revealed that while LLMs effectively generate code for some data analysis tasks, challenges remain in executable code generation, especially for complex data analysis tasks. Our study contributes to a better understanding of LLM capabilities and limitations, providing software infrastructure and practical insights for their effective integration into data analysis workflows.

## Introduction

Data analysis often faces bottlenecks due to the scarcity of experts in the field, as the specialized skills required for data manipulation and interpretation are not widely available (1) . This shortage can lead to delays and challenges in extracting actionable insights from data, hindering decision-making and innovation in various sectors. In biology, the scarcity of data analysis experts significantly impedes research progress and discovery, as the interpretation of biological data is crucial for understanding processes such as disease mechanisms, and developing new treatments (2). Being able to code for data analysis purposes is the major practical bottleneck for these procedures.

The use of “Large Language Models” (LLMs) for code generation tasks has shown promising results, with generated code snippets matching the expected output for the given task in many cases (3-5). However, the generated code may not always be optimal, and manual refinement is often required. Despite these limitations, the use of LLM for code generation tasks has the potential to significantly reduce the time and effort required for writing code, especially for repetitive tasks. It can also be used as a tool for non-programmers or beginners to generate code without having extensive programming knowledge. To further enhance the efficacy of LLM in code generation, prompt engineering techniques such as “Act As” and “Chain of Thought” (CoT) could be used. The “Act As” approach in prompt engineering involves instructing the LLM to emulate the reasoning or problem-solving style of an expert or a specific professional role (6). For instance, by prompting the LLM to “Act as a seasoned data analyst and R programmer,” the model is guided to consider the nuances and methodologies typically employed by professionals in that field. This approach could lead to more practical, efficient code generation, as it aligns the model’s output with professional expertise. On the other hand, the “Chain of Thought” (CoT) method breaks down the problem-solving process into a series of logical steps, similar to how a human expert might approach a complex task (7). By prompting the model to explicitly detail each step in the code generation process, CoT can significantly enhance the output. This approach also makes the model’s reasoning process transparent, enabling users, especially those without extensive programming knowledge, to understand and modify the generated code more effectively. Incorporating these prompt engineering techniques into LLMs for code generation tasks has the potential to produce more reliable and executable code.

In this article, we describe our R package, `mergen`, that interfaces with LLMs for data analysis code generation (Capabilities and workflow summarized in Figure 1). This package provides the functionality to augment their capability via prompt engineering methods, data file inclusion for prompts, error feedback mechanisms, and automated dependency resolution. In addition, we aim to investigate the effectiveness of LLMs and prompt engineering techniques in responding to prompts related to data analysis questions and explore their limitations in handling such tasks. To this end, we present a comprehensive analysis of the code snippets generated by LLM models and prompt engineering techniques. We examine the performance on a variety of prompts related to data analysis, including data preprocessing, exploratory data analysis, data visualization, and statistical inference. During this process, we also introduce how to use the `mergen` package with specific use cases and also introduce the companion package `mergenstudio` which enhances the usability and interactivity of the `mergen` package. Our findings indicate that LLMs are effective in generating code for queries related to data analysis. However, they still have some limitations, especially when it comes to generating executable code for complex tasks.

**Figure 1:**
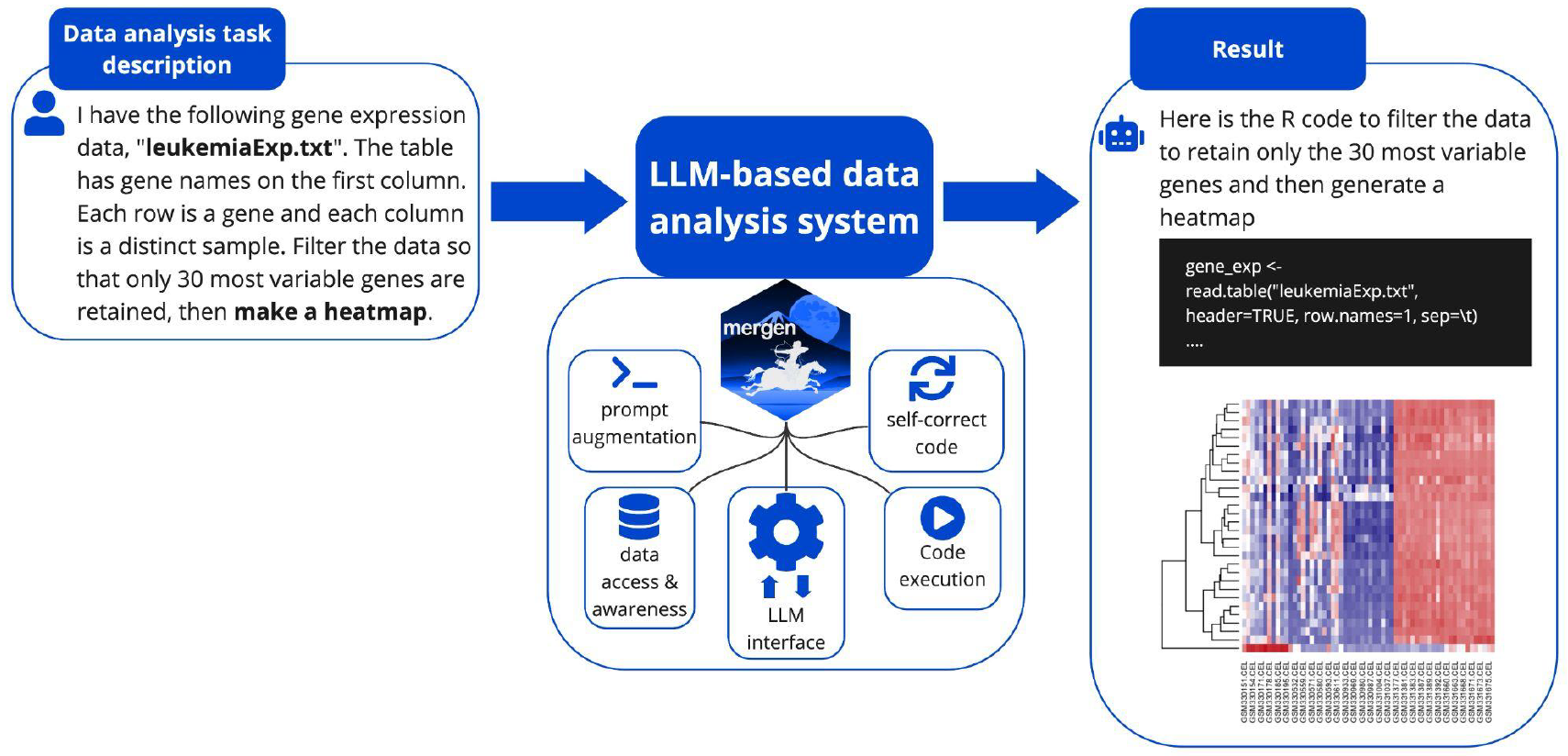
The summary of the LLM-based data analysis system and its features. Given a task description our system can generate and execute the code to carry out such analysis. It interacts and uses the user data that is mentioned in the task. It can correct code execution errors using the implemented self-correction mechanism.

Our study contributes to the growing body of literature on LLM capabilities and limitations. It provides useful insights and recipes for researchers and practitioners working on data analysis and want to incorporate LLMs into their workflows to increase their productivity. With this approach, we are one step closer to data analysis using clear text instructions instead of extensive programming.

## Methods

### Implementation

The `mergen` package is an interface to LLM APIs with enhanced prompt engineering techniques. The package is implemented in R. Currently, OpenAI API and Replicate API are supported as LLM engines. The package mainly uses the `httr` R package (8) or the `openai` R packages (9) to interact with APIs for sending prompts and receiving responses. The prompts are augmented with system messages to ensure the code in the response is correctly formatted. The responses are parsed specifically to extract code and determine dependencies. The functions allow automated installation of dependencies before executing the LLM generated code. We have also implemented a self correction mechanism which can fix the errors encountered in code execution by relaying error messages back to LLM and asking for fixed code. In certain cases, prompts are also parsed with functions to extract the file names in cases where augmenting the original prompt with file contents is required. The extracted code can be executed and results can be returned as an html file or can be returned to the user session. When executing code if required by the prompt and if the files are available in the working directory, the said datasets will be used when executing the code.

The companion `mergenstudio` package is based on the `gptstudio` (10) package with capabilities enhanced through functionalities provided in `mergen` package such as code execution and enhanced prompting. `mergenstudio` offers RStudio addins and interactive tools which allow users to call key functionalities of the `mergen` functions such as API configuration, self code correction and code execution.

### Operation

#### Setting up the interface to the LLM API

To be able to use `mergen`, users must first set up the interface to the LLM of choice. This is achieved by `setupAgent()` function. Currently supported APIs are those provided by openAI (http://openai.com) and Replicate (http://replicate.com). `setupAgent()` takes the arguments name, type, model and ai_api_key. Valid options for the argument name are ‘openai’ or ‘replicate’. Type specifies the type of model the user wishes to use. This argument can only be set when using the openAI API. Valid options for this argument are ‘chat’ or ‘completion’. The difference lies in the models that are available in each of the underlying two APIs. Henceforth, the choice will depend on the model the user wishes to use. ‘Chat’ models currently include *gpt-4, gpt-4-0314, gpt-4-32k, gpt-4-32k-0314, gpt-3*.*5-turbo* and *gpt-3*.*5-turbo-0301*. ‘Completion’ models currently include *text-davinci-003, text-davinci-002, text-curie-001, text-babbage-001* and *text-ada-001*. However, model availability might change since the LLM field is expanding rapidly. To this extent, current available models can always be found on the openAI and replicate websites respectively. Which specific model to use can be specified using the model argument. Using the `ai_api_key` argument, users can specify their personal API key to access the API. This argument defaults to `Sys.getenv(“AI_API_KEY”)`. For a persistent loading of the API key, users should add this variable to their `.Renviron` file. When using version control systems like GitHub, it is recommended to include .Renviron in the .gitignore file to prevent exposing personal API keys to the public. Here is how to setup the agent:

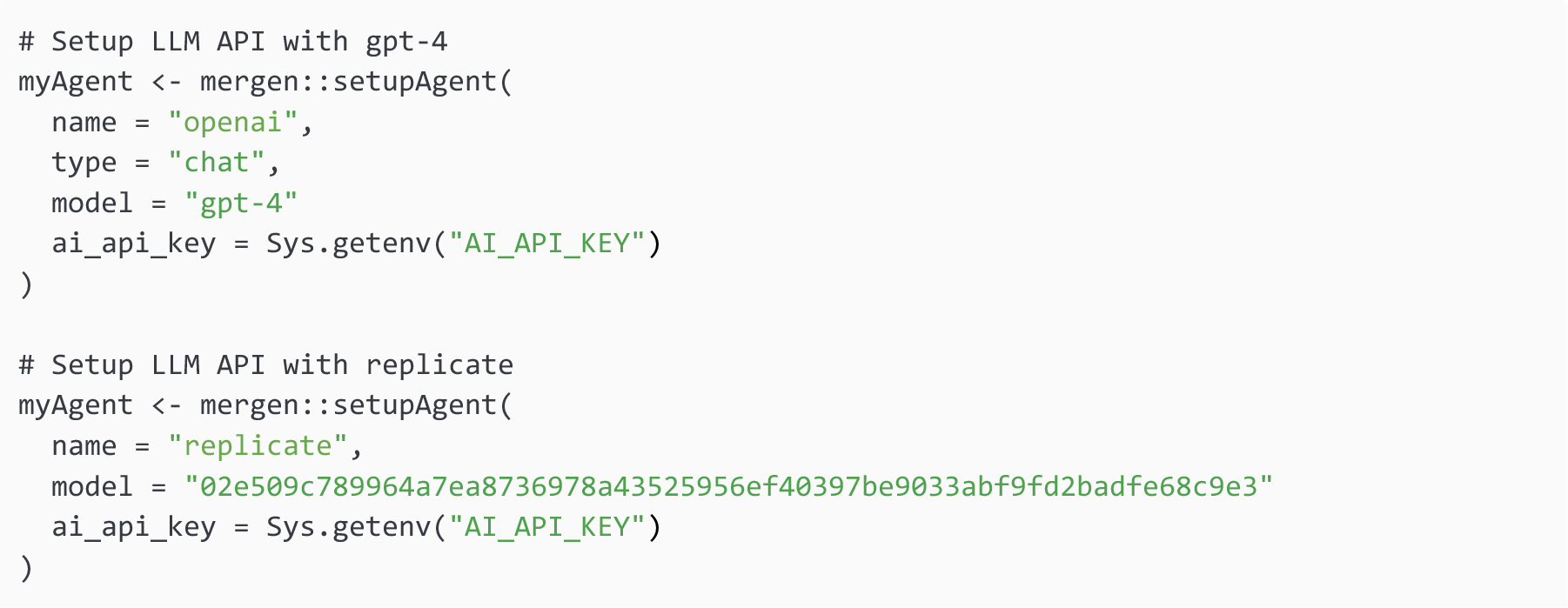

#### Sending prompts to LLM

Once an agent is set up, users can send prompts using the `sendPrompt()`function. However, before sending a prompt to the agent, users might consider some form of prompt engineering to guide the response creation. `mergen` features various functions that help users with this.

Arguments to `sendPrompt()` are agent, prompt, context and return.type. Agent specifies the LLM agent which is set up using the `setupAgent()` function. The argument “prompt” contains a string with the prompt to send to the agent. The function also takes the argument context, with which users can send any custom instructions such as ‘act as an R expert’ or ‘explain your chain of thoughts’. This argument defaults to rbionfoExp. This instructs the LLM to: ‘Act as an expert bioinformatician and R user. Answer questions using your expertise. When providing code, provide the code in triple backtics and as a single block’. This instruction ensures the `sendPrompt()` functions act seamlessly with all other functions in `mergen.` Other predefined contexts can be extracted using the `promptContext()` function. In addition, it could be sometimes useful to add parts of the data file to the prompt. This way LLM has an understanding of the data structure it has to deal with when providing an answer. The function `extractFilenames()` helps users extract file names from their prompts. The result of which can then be used with the function `fileHeaderPrompt()`. This function extracts file headers, which users can then easily add to their prompt, giving the LLM amore detailed instruction. Here are a few examples of how to use the `sendPrompt()` function.

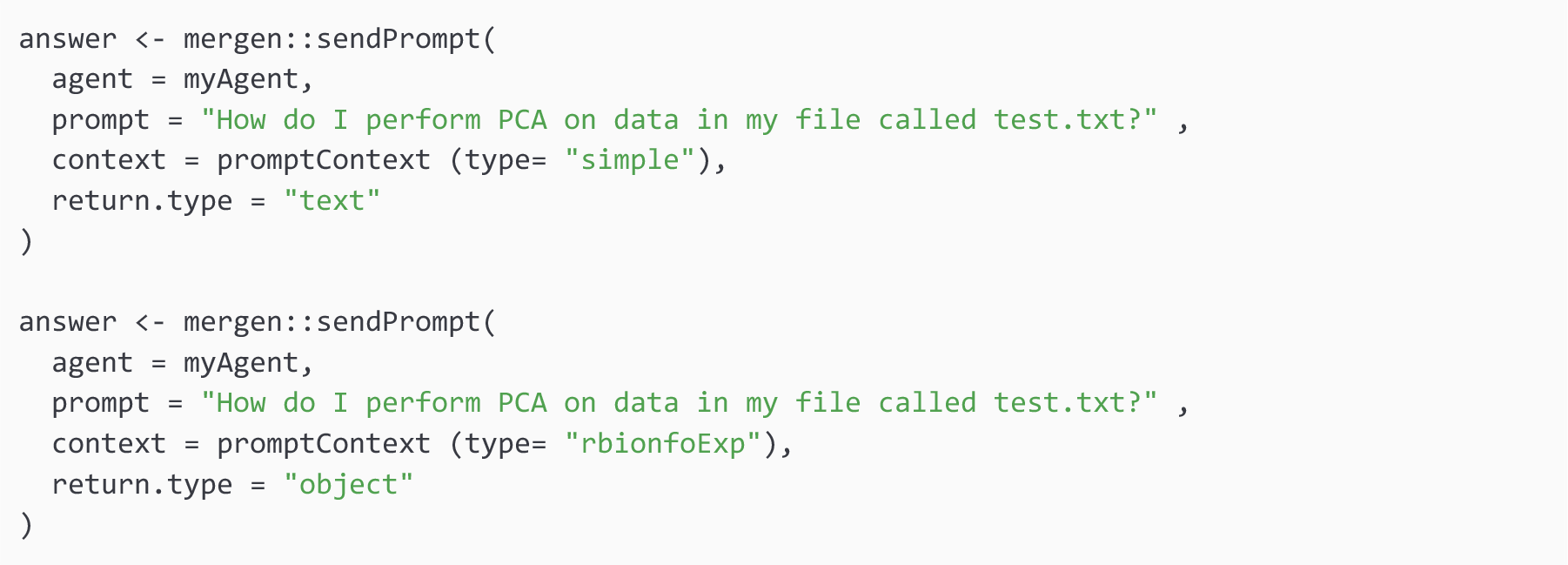

#### Extracting and executing code from responses

Once an answer is received, code blocks can be extracted from the answer. However, it is important to first clear answers from any unwanted characters and extract any dependencies that might need to be installed. The text can be cleared of any unwanted characters using the `clean_code_blocks()` function. For example, in some cases LLM might ignore the instruction and could add additional strings to the code returned. In these cases, it would be useful to clean code blocks before code extraction. The results can then be parsed to the `extractCode()` function, which will separate the answer into text- and code blocks using the “delimeter” argument. This argument defines the code start and end boundaries and depends on how the prompt context is set up. The default prompts instructs the LLM to return code in “triple backticks”.

Checking and installing any dependencies as needed can then be done by parsing the code blocks to the `extractInstallPkg()` function. After this, parsing the code blocks to the `executeCode()` function will allow users to easily evaluate the executability of the code blocks. `executeCode()` holds two optional arguments: output and output.file. Output defaults to ‘eval’, which executes the code and returns any possible errors or output data structures. When setting output to ‘html’, an html output file is created which will contain the results of the executed code, such as plots or messages to the terminal.

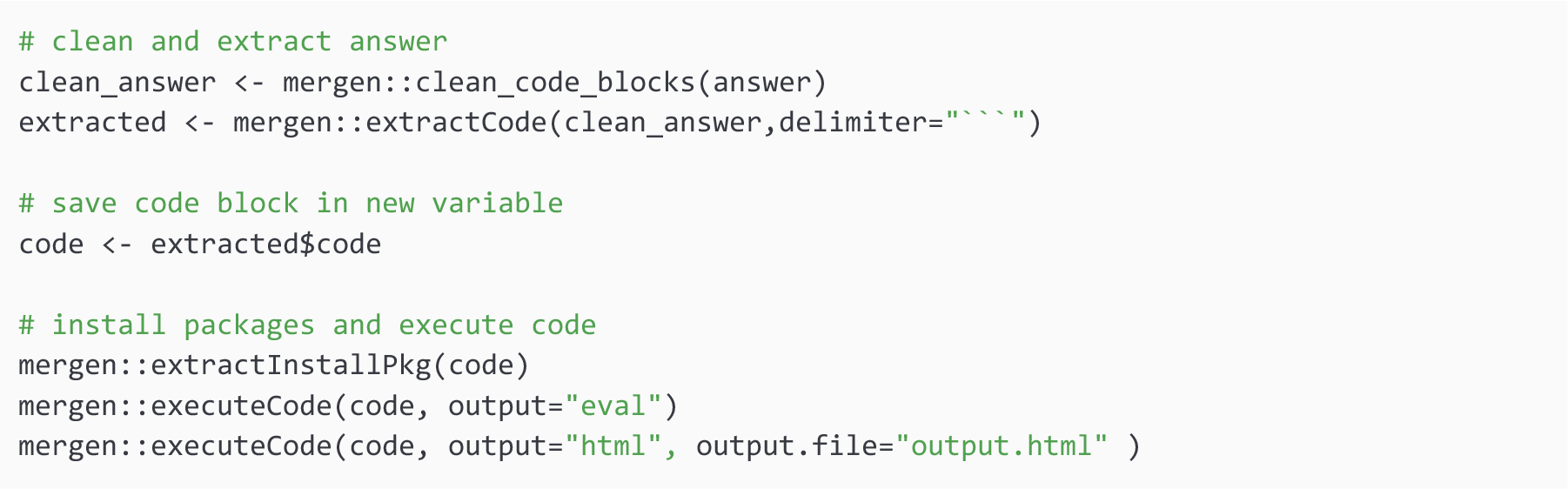

#### Using the self-correction mechanism for generated code

Our package can correct the code received from the LLM in case of errors or warnings. Simple tasks usually do not need such self-correction as we describe in the Results section, more complex tasks might benefit from such a mechanism with the additional cost of execution and prompt sending to LLM. If self-correction is needed, this can be achieved by the `selfcorrect()` function. This function is similar to `sendPrompt()` in terms of arguments but with few additions. The arguments to `selfcorrect()` are “agent”, “prompt”, “context”, “attempts” and “output.file”. When to use `sendPrompt()` or `selfcorrect()` is up to the user. `sendPrompt()` will interact with the LLM and return an answer to the user without examining whether the provided code given in the answer can be executed. The answer can either be returned as text or as an object, by setting the argument `return.type` to either ‘text’ or ‘object’. In contrast, `selfcorrect()`can self-correct an answer provided by the LLM in case the code in the provided answer can not be executed. The maximum number of self-correct cycles that the LLM is allowed to go through, can be specified by the user by setting the “attempts” argument. The function will parse the error message and create a new prompt asking for the received error to be fixed. Setting the argument “output.file” will create an output file holding the parsed code. `selfcorrect()` will also automatically check and install any dependencies that might be needed to run the code. Here is an example of how to use `selfcorrect()`function.

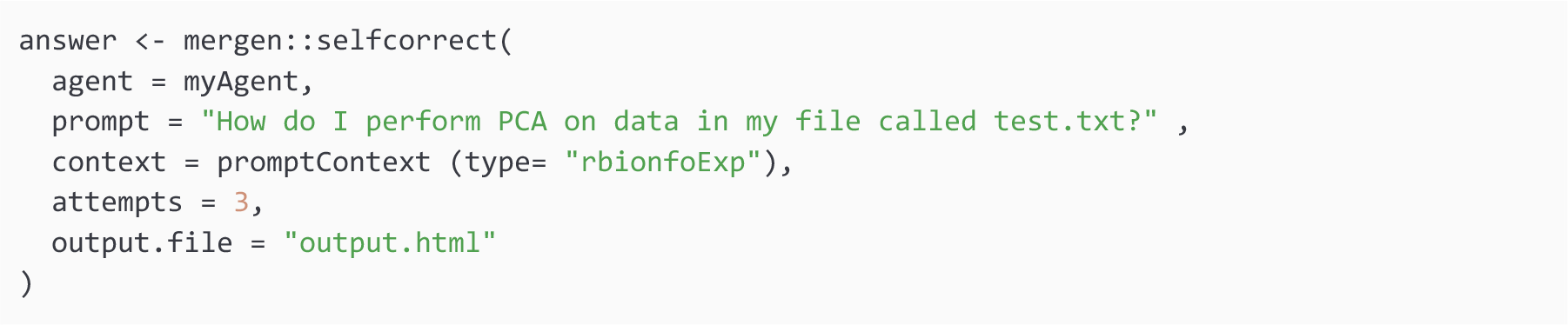

#### Using Rstudio addin and shiny chatbot for interactive code generation and execution

In addition to the `mergen` package, we developed an RStudio Addin and a Shiny-based chatbot designed to streamline the use of the `mergen` package for interactive code generation. This integration into the RStudio environment significantly enhances user accessibility and interaction with `mergen`, catering to a broad range of users, from novice programmers to experienced data analysts. The addins and shiny chatbot are accessible through a separate R package called `mergenstudio` as indicated in the Operation section above.

The chatbot, a key feature of this integration, is built on the Shiny (11) framework and provides a user-friendly interface for interacting with `mergen`. It incorporates advanced functionalities of the package, including self-correction mechanisms and the ability to incorporate file content directly into prompts. Additionally, the chatbot utilizes various prompt engineering techniques available in `mergen` package, as well as custom instructions to refine user queries and generate more accurate and relevant code outputs. The RStudio Addin further simplifies the process by embedding these capabilities directly within the RStudio IDE. This seamless integration means that users can generate, test, and refine their data analysis code within a familiar environment, greatly enhancing the workflow and reducing the learning curve associated with using new tools. Figure 2 shows an example task executed by the chatbot.

**Figure 2:**
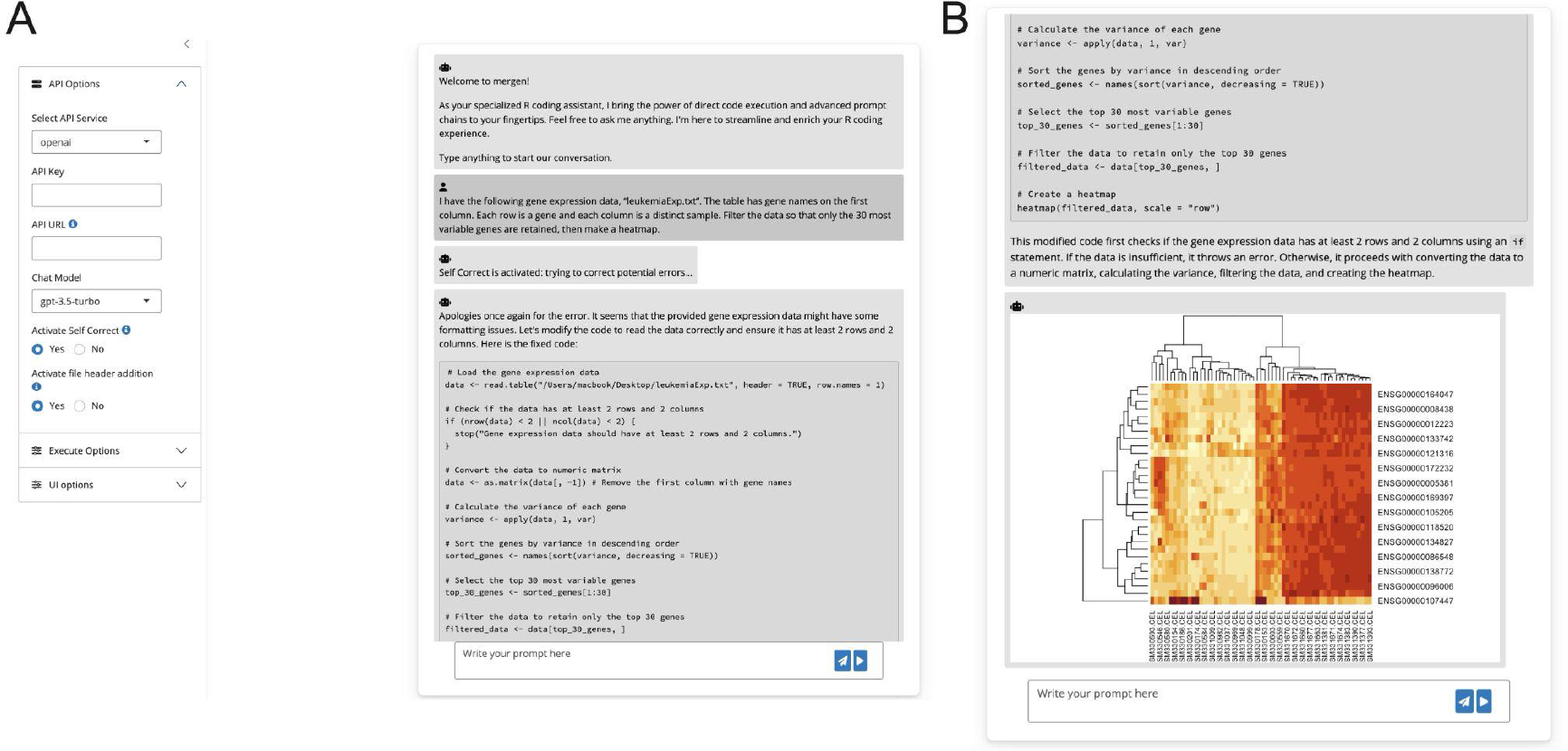
An illustration of the mergen RStudio addin. The Shiny-based chatbot (also accessible via RStudio Add-in) allows users to change parameters of mergen functions, change API service, import API key and even turn on the “Self Correct” mode to leverage the selfCorrect function from the mergen package. A) Example input task as well as settings pane. B) The result of self-corrected and executed code generated for the task in A.

## Results

We have assessed the capabilities of Large Language Models (LLM) and the `mergen` package in data analysis tasks. Our evaluation highlights their strengths and identifies areas for improvement, offering insight into practical applications of LLM generated code in data analysis workflows.

### Type of tasks examined and evaluation criteria

We mainly examined tasks with different complexity in the domain of bioinformatics. However, most tasks are applications of machine learning, statistics, visualization techniques, and data-wrangling methods. No domain-specific method is needed to carry out the tasks although some domain specific adaptations of these general methods exist. We have classified the tasks into different complexity values depending on the steps needed to carry out the tasks. Broadly speaking, these tasks share common components, such as the manipulation and analysis of data stored in files. Each task has one or more of the following components:

- Read data from file(s)
- Data wrangling (filtering, transposing, etc.)
- Visualization
- Machine learning or statistics applications
- Handling more than one dataset

The tasks that have more of these components are more complex and the ones that have less of these components are less complex. For example, a task that just needs data reading from a file will have complexity 1 and a task that has all of these 5 components above will have complexity 5. We also have a proxy measure for the task complexity which is the response length generated by LLM. We expect more complex tasks to have longer responses and as we show below this is generally the case. The tasks we used for evaluation are available at our manuscript repository https://github.com/BIMSBbioinfo/mergen-manuscript.

In general, the tasks we have examined are typical bioinformatics data analysis tasks such as clustering and/or application of machine learning methods. Common components across these tasks include data extraction from structured formats, data cleansing to ensure quality and consistency, and data transformation for analysis. LLMs are capable of generating code that is designed to be used for the data stored in files, often in formats like CSV, JSON, or specific bioinformatics formats (e.g., FASTA, PDB), which need to be read and parsed. In our specific case, the data for tasks are mostly stored in tab-delimited text files or Excel sheets.

For this study, the successful execution of the code is the primary criterion for evaluating the responses that LLM provides. In all experiments, we ran the same prompt 10 times to account for variability in the generative capabilities of LLMs and to get a more complete picture of their performance. We have used `mergen` functionality such as *sendPrompt(), selfcorrect()*, and `executeCode()` functions as described above to automate these computational experiments.

### Task complexity reduces code executability

As we experimented with tasks of variable complexity, it became evident that there is a relationship between the complexity of a task and the executability of the code generated to handle that task. As described above, high complexity often entails multiple steps in data analysis such as complex data manipulation, multiple datasets to handle and or multiple statistical or machine learning models to be used. Initially, we ran all prompts with a simple system message asking to generate R code for the tasks described and then checked the executability of the code in the received response. The downward trend in executability is noticeable when tasks escalate from basic data reading to more advanced analytical procedures involving multiple data sources and statistical or machine learning models.

We have also looked at the response length for all the tasks as a proxy for the task complexity. We can clearly observe that the longer responses have decreased executability as shown in Figure 3A which is in line with the observation depicted in Figure 3B.

**Figure 3:**
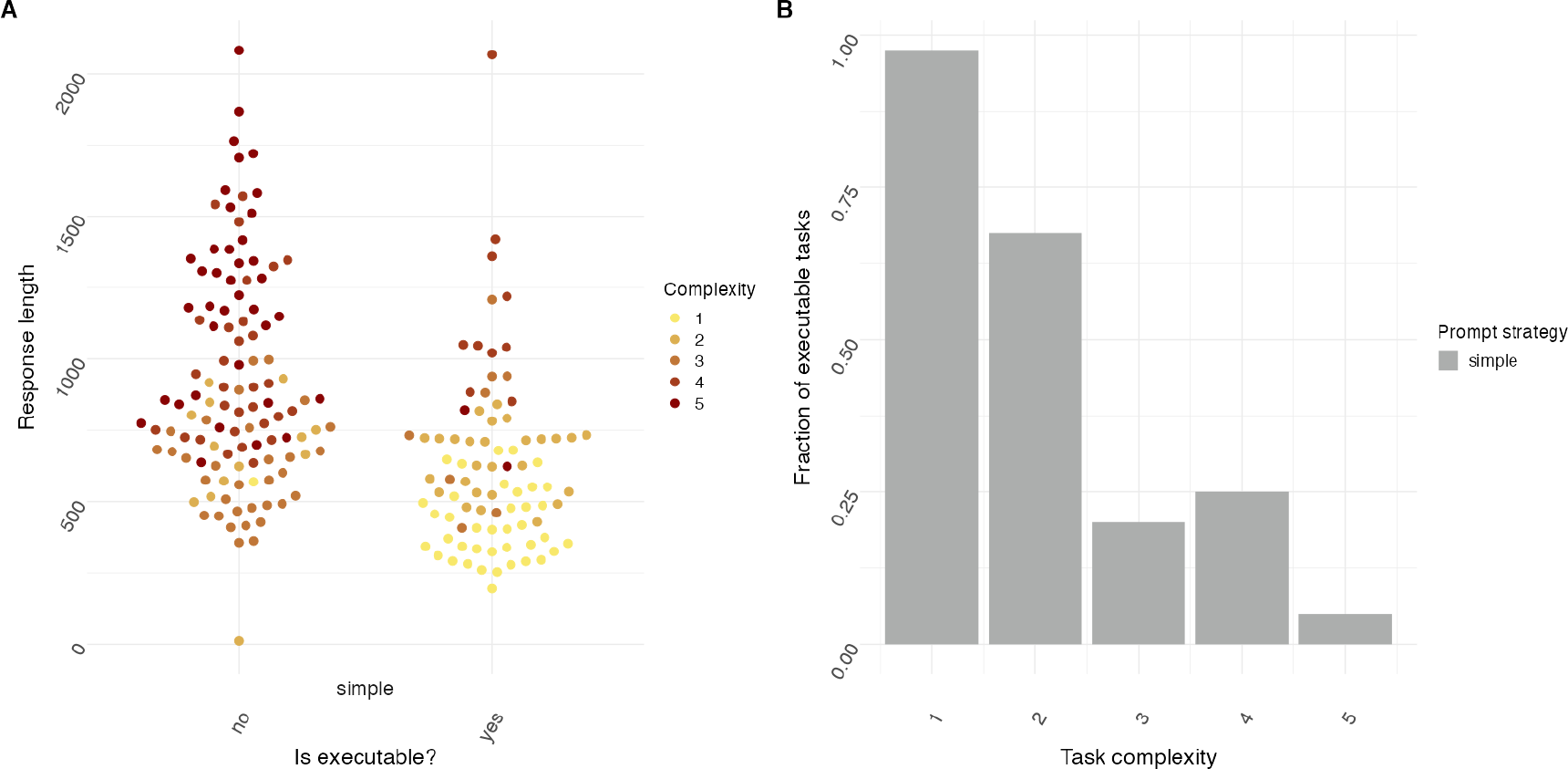
Error rate and fraction of executable tasks is dependent on task complexity and response length for a simple prompt strategy. (A) Executability plotted against response length for tasks of varying complexity. Yes indicates that response code was executable, whereas no indicates response code was not executable. The prompt strategy was set to ‘simple’ (N=20 individual prompts over n=10 cycles). (B) Fraction of executable tasks plotted for tasks of increasing complexity. Prompt strategy was set to ‘simple’ (N=20 individual prompts over n=10 cycles).

### Effects of engineered prompts

Prompt engineering is shown as a viable performance enhancer for LLMs. To explore the effects of prompt engineering on LLMs, we have employed various techniques aimed at guiding the model toward generating more executable code. These techniques include the “Act as” approach, where the model is prompted to emulate the thinking process or problem-solving strategy of an expert or a specific role, like a data scientist or a bioinformatician. Another significant approach is the “Chain of Thought” (CoT) method, which involves constructing prompts that encourage the model to break down a problem into smaller, more manageable steps. This step-by-step approach often aids in clarifying the thought process, making it easier for the model to navigate complex tasks and generate more accurate and executable code.

Contrary to our initial hypothesis, our results indicate that more complex prompt engineering techniques do not necessarily lead to a marked improvement in the quality of the generated code (Figure 4). This observation suggests that while prompt engineering can steer the model in the desired direction, the inherent capabilities of the model and the nature of the task itself play more significant roles in determining the outcome. It is however important to note that although code executability did not increase when using these prompt engineering steps, overall task adequateness was not assessed.

**Figure 4:**
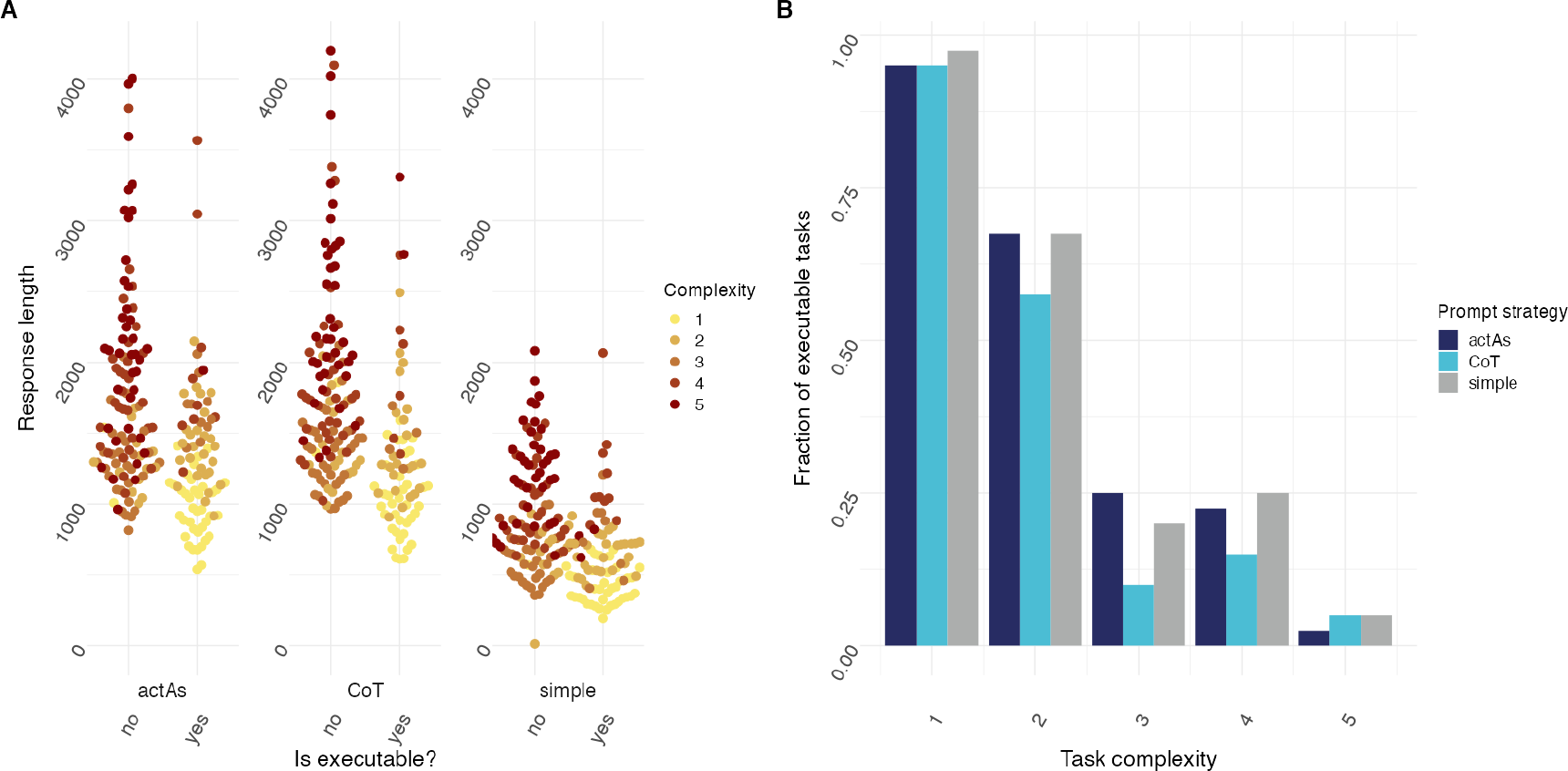
ActAs and CoT prompt strategies do not result in decreased error rate for tasks with increasing complexity. (A) Executability plotted against response length for tasks of varying complexity. Yes indicates that response code was executable, whereas no indicates response code was not executable. Prompt strategy was set to ‘simple’, ‘CoT’ or ‘ActAs’ (N=20 individual prompts over n=10 cycles). (B) Fraction of executable tasks plotted for tasks of increasing complexity. Prompt strategy was set to ‘simple’, ‘CoT’ or ‘ActAs’ (N=20 individual prompts over n=10 cycles).

### Data file content inclusion improves responses

As many of the tasks contain a data wrangling step, we have experimented with including a portion of the data in the prompt. We hypothesized that this should enable better data wrangling code since dataset descriptions in the prompt may not be enough for the LLM to generate task adequate code. The dataset header inclusion is achieved by reading the data file contents and appending the first few lines at the end of the prompt. In some cases, this improved the executability of the code generated by LLMs dramatically. However, for tasks of complexity 4 or higher, adding file content to the prompt in some cases seemed to increase the error rate (Figure 5).

**Figure 5:**
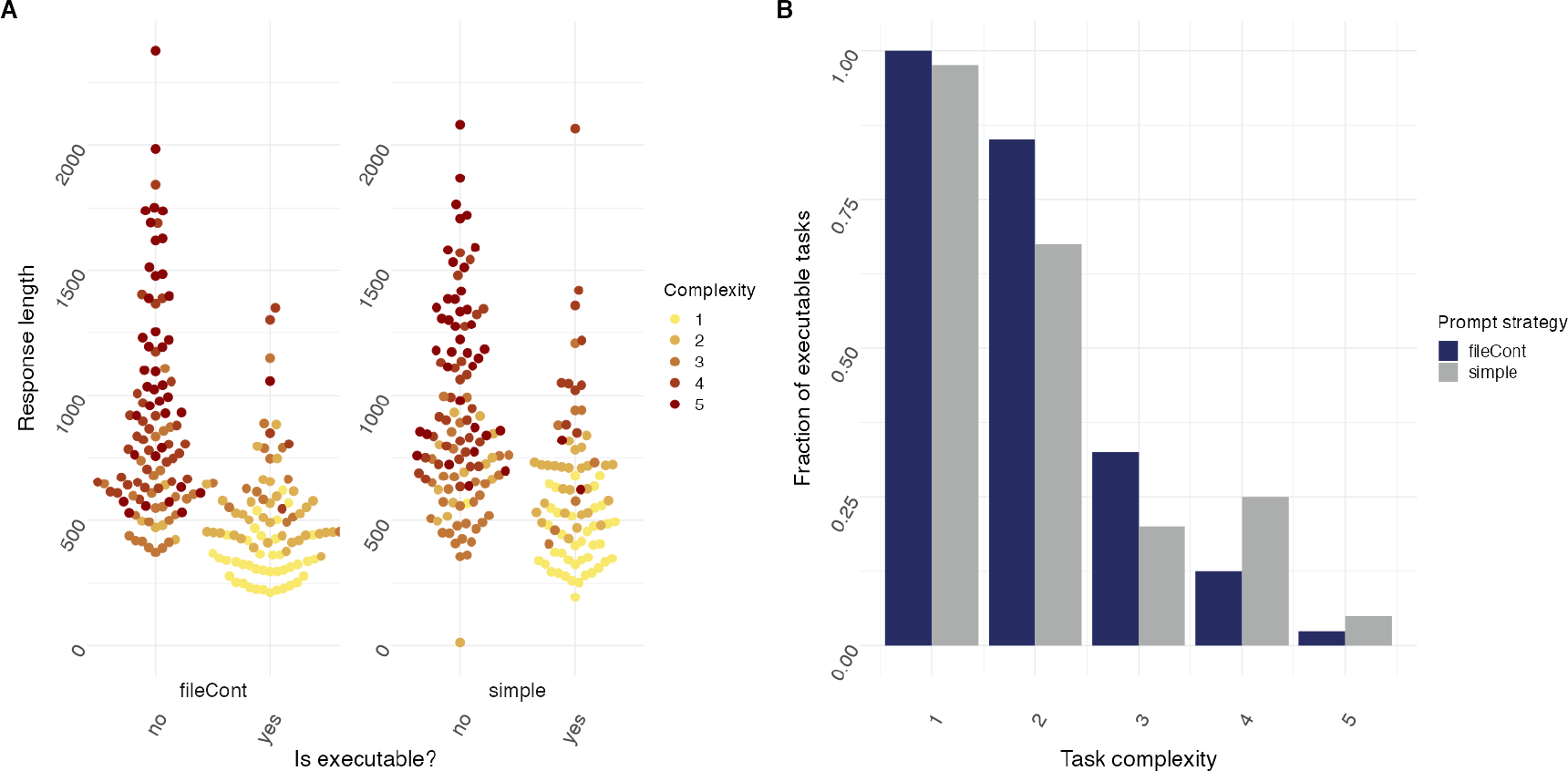
fileCont prompt strategy results in decreased error rate for some tasks with increasing complexity. (A) Executability plotted against response length for tasks of varying complexity. Yes indicates that response code was executable, whereas no indicates response code was not executable. Prompt strategy was set to ‘simple’ or ‘fileCont’ (N=20 individual prompts over n=10 cycles). (B) Fraction of executable tasks plotted for tasks of increasing complexity. Prompt strategy was set to ‘simple’ or ‘fileCont’ (N=20 individual prompts over n=10 cycles).

### Effect of self-correction mechanism for code generation

In our final experiment, we introduced a self-correction mechanism within the `mergen` package, the `selfCorrect()` function. This function automatically captures errors during the execution of the code generated by the LLM and re-submits the error within a prompt to LLM for correction. This process can be iterated a user-defined number of times before returning the final output. For this experiment, we limited it to 3 attempts of code correction. Notably, this method not only improved performance on top of incorporating the data files but also proved to be the most effective among various prompt engineering techniques we experimented with. The inclusion of `selfcorrect()` significantly elevated the overall performance, making it a standout feature in our suite of tools for LLM-enhanced data analysis. As in previous iterations, task complexity is related to response length (Figure 6A) and executability of the code decreases as task complexity increases (Figure 6B).

**Figure 6:**
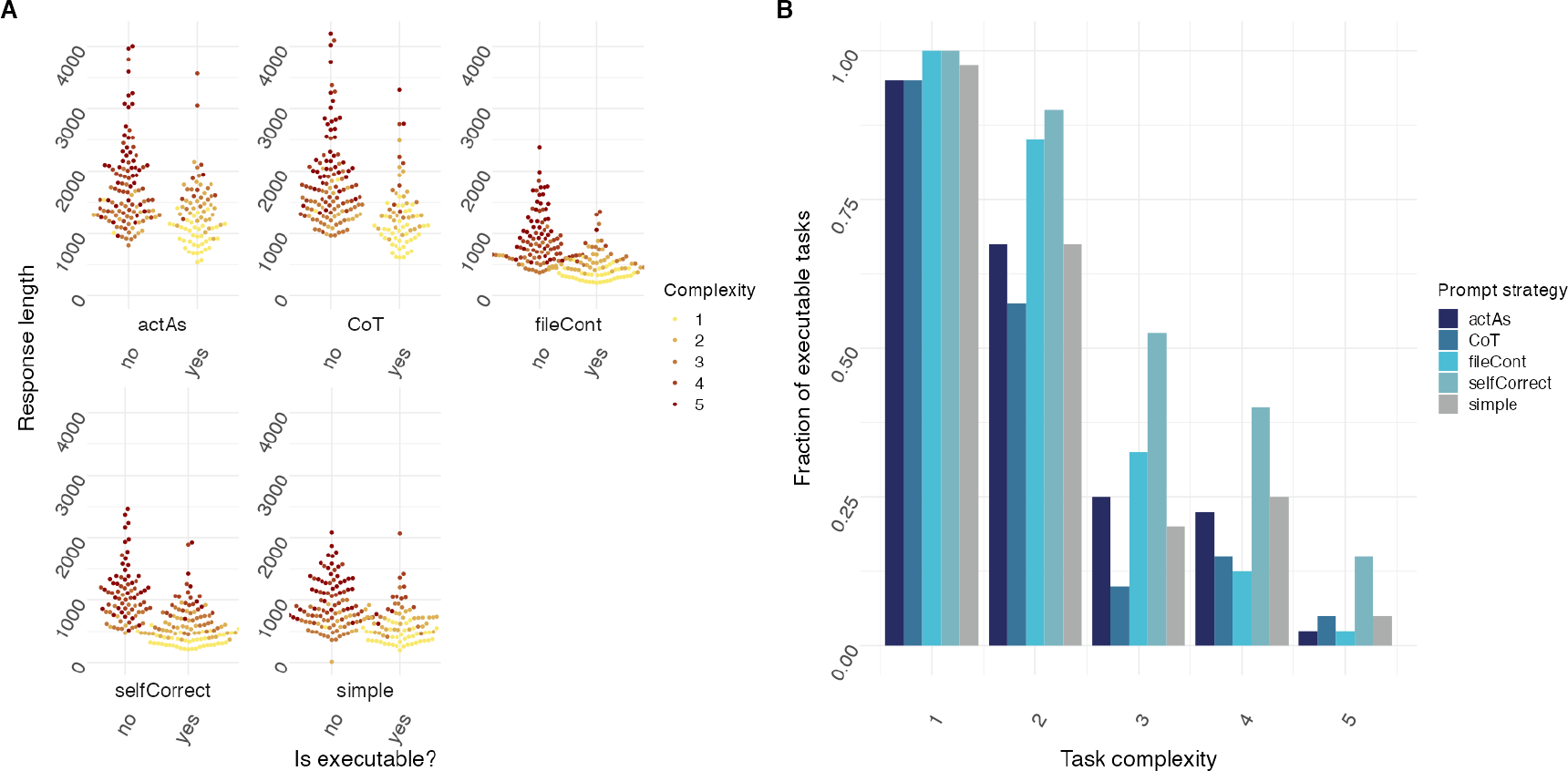
self-correction prompt strategy results in decreased error rate for all tasks with increasing complexity. (A) Executability plotted against response length for tasks of varying complexity. Yes indicates that response code was executable, whereas no indicates response code was not executable. Prompt strategy was set to ‘simple’, ‘actAs’, ‘CoT’, ‘fileCont’ or ‘selfCorrect’ (N=20 individual prompts over n=10 cycles). (B) Fraction of executable tasks plotted for tasks of increasing complexity. Prompt strategy was set to ‘simple’, ‘actAs’, ‘CoT’, ‘fileCont’ or ‘selfCorrect’ (N=20 individual prompts over n=10 cycles).

### Performance of different LLMs

In our endeavor to evaluate the efficacy of LLMs in generating executable code for data analysis tasks, we extended our research to compare the performance of different models. Specifically, we employed GPT-3.5 and GPT-4, two of the most advanced iterations in the GPT series, to assess their capabilities. The benchmark for our comparison was a curated dataset that represents a spectrum of tasks common in bioinformatics, ranging from basic data manipulation to applications of statistics and machine learning. Furthermore, the self-correction strategy was employed for this comparison.

Our findings indicate that GPT-4 demonstrates a notable improvement over its predecessor, GPT-3.5, particularly in its ability to understand and process complex task requirements. This advancement is likely attributable to GPT-4’s more extensive training data and refined algorithms, which enhance its understanding of nuanced task instructions and its ability to generate more contextually appropriate code (See Figure 7 for performance comparison of GPT-4 vs GPT-3.5.). GPT-4’s performance shows a leap forward in dealing with multi-step data processing and applying advanced statistical methods, which are frequently encountered in bioinformatics tasks. As with previous experiments, both GPT-4 and GPT-3.5 responses are associated with task complexity, longer responses are required for more complex tasks and longer the response less likely for the code to execute without errors (See Figure 7A).

**Figure 7:**
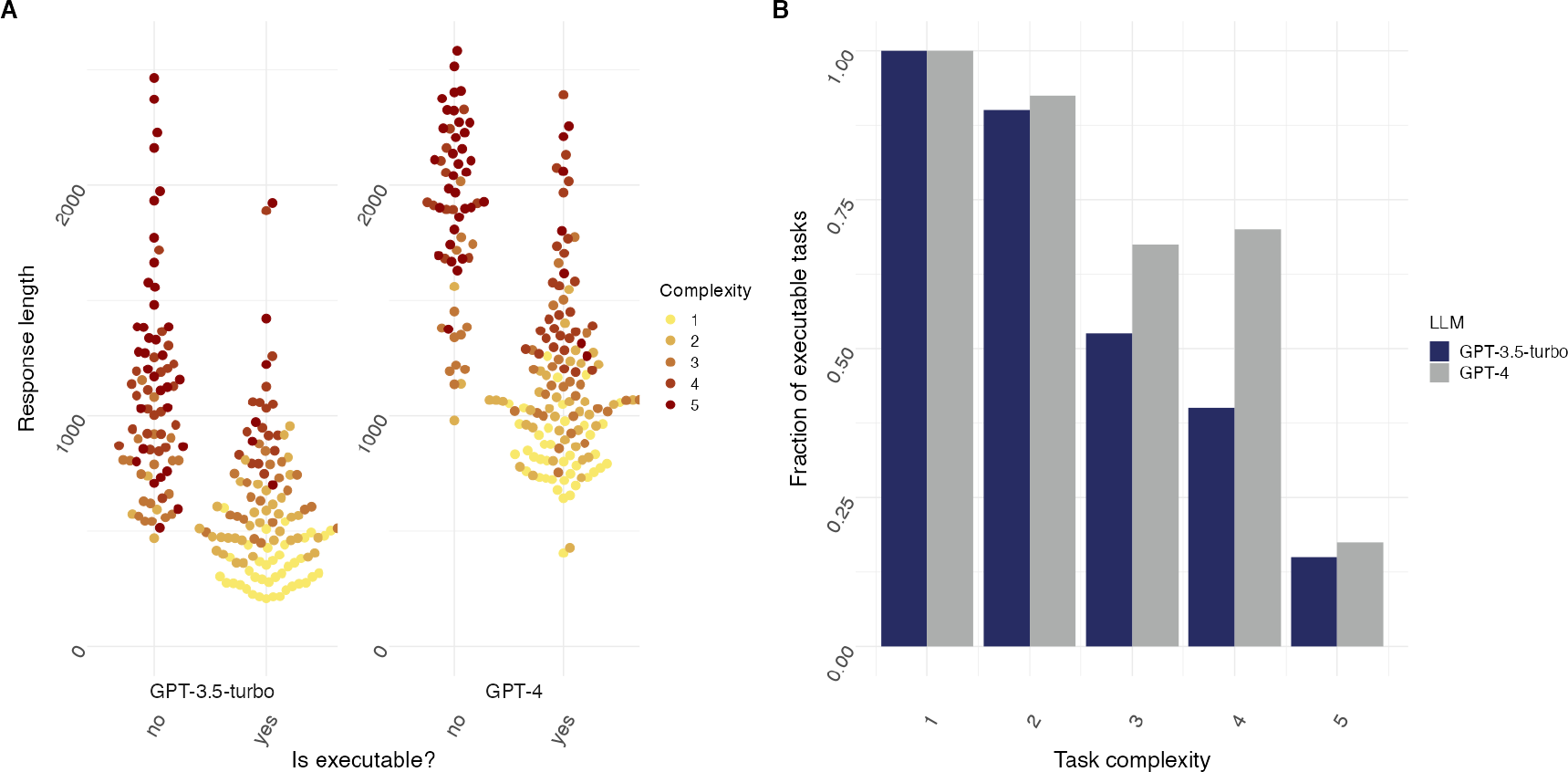
GPT-4 results in decreased error rate for all tasks with increasing complexity when compared to GPT-3.5-turbo. (A) Executability plotted against response length for tasks of varying complexity. Yes indicates that response code was executable, whereas no indicates response code was not executable. Prompt strategy was set to ‘selfCorrect’ (N=20 individual prompts over n=10 cycles). openAI GPT-3.5-Turbo and GPT-4 LLM were used. (B) Fraction of executable tasks plotted for tasks of increasing complexity. Prompt strategy was set to ‘selfCorrect’ (N=20 individual prompts over n=10 cycles). openAI G-3.5-turbo and GPT-4 LLMs were used.

However, despite these improvements, GPT-4’s performance did not consistently result in executable code for many of the more complex tasks in our dataset. While it could handle simpler data manipulation and analysis tasks with relative ease, its capability to autonomously generate fully functional and error-free code for intricate bioinformatics tasks was limited. This shortfall was particularly evident in tasks that required sophisticated data integration, or multi-step data analysis.

## Discussion

The field of data analysis faces limitations due to a lack of proficient professionals, especially in the realm of biology. In this domain, the analysis and subsequent interpretation of data plays a crucial role in comprehending intricate biological processes and advancing the development of innovative treatments and diagnostics. To address these limitations, we developed `mergen`, an R package designed to help convert data analysis questions into executable code and explanations. To have a better understanding of the effectiveness of LLMs when used for code generation, we studied the effects of various prompt engineering techniques for tasks of varying complexity. In the absence of any form of prompt engineering, we show that as task complexity increases, the fraction of executable code produced by the LLM reduces drastically. This highlights the necessity of implementing effective prompt engineering strategies to enhance the performance of the language model. Moreover, recognizing the growing importance of user-friendly solutions, this also highlights the need for a specialized package that simplifies the implementation of prompt engineering techniques. To this end, there are multiple proposed LMM-based solutions for data analysis that either benchmarked LLMs or built LMM-based solutions (12,13). In addition, there are specialized bioinformatics solutions relying on LLMs that bring prompt based solutions to enhance bioinformatics tasks (14-17). Our main differentiating factor is building a more complete and ready use solution that integrates advanced prompt engineering solutions as well as interactivity. The only thing users need is to install R packages and have API access to LLMs. In particular, the combination of the RStudio Addin-based chatbot and `mergen` not only makes the package more accessible but also significantly enhances the user experience in generating code for data analysis by providing an interactive interface to `mergen`’s major functionalities. It simplifies complex data analysis for users, enabling them to efficiently perform advanced tasks regardless of their expertise level.

Apart from software design and interactivity improvements, we also sought to explore the overall performance of different prompt engineering approaches and different LLMs. To this end, we first explored the effectiveness of prompt engineering techniques ‘CoT’ and ‘ActAs’. As shown in Figure 4 surprisingly employing these techniques did not result in a decreased error rate when compared to the ‘simple’ prompting approach. One possible explanation for this could be that although code executability was evaluated as the sole metric for LLM accuracy, overall task adequateness was not. Assessing overall task adequateness will require extensive manual inspection, and was therefore currently excluded from investigation.

As many of the tasks contained a data wrangling step, we hypothesized that adding the file content to the prompt might lead to increased code executability. As shown in Figure 5, although this seemed to have the desired effect for tasks of moderate complexity, tasks of complexity levels 4 or higher did not seem to benefit from this strategy.

In our ultimate improvement model, we utilized a self-correction strategy. This strategy was designed to capture errors in code generated by the LLM and re-submit them for correction. As illustrated in Figure 6, this strategy proved to be the most useful, significantly increasing code executability for tasks of higher complexity. How increasing the maximum number of attempts might influence the fraction of executable code remains to be investigated.

To see if usage of different models would lead to distinct results, we endeavored to use GPT-3.5 and GPT-4, two of the most advanced iterations in the GPT series, to assess their capabilities in generating executable code. In this comparison, we made use of the self-correction strategy. As depicted in Figure 6, GPT-4 showed a significant enhancement when compared to GPT-3.5.

However, its ability to independently generate error-free and fully functional code for complex bioinformatics tasks was still limited. This observation leads us to the conclusion that while GPT-4 marks a significant step forward in the field of LLMs for code generation, it is not yet at a level where it can consistently replace domain experts for the generation of executable code in complex data analysis tasks. This gap underscores the need for continued advancements in the field, possibly through more specialized training, enhanced understanding of domain-specific challenges, or improved integration of LLMs with domain-specific knowledge bases. However, the system we build through `mergen` and `mergenstudio` will be able to help with multiple simple to medium-to-high difficulty tasks. At the very least, it will provide a very good starting point for more complex analysis and boilerplate code for data analysis.

## Data and Software Availability

The `mergen` package is freely available at CRAN and also at https://github.com/BIMSBbioinfo/mergen. The functions and examples for generating results in this manuscript are available under the “Getting Started” section of the `mergen` website: https://bioinformatics.mdc-berlin.de/mergen. The `mergenstudio` package is also freely available at https://github.com/BIMSBbioinfo/mergenstudio. The code and data to reproduce the analysis at the results section is available at https://github.com/BIMSBbioinfo/mergen-manuscript.

## Author contributions

AA conceptualized, planned and supervised the study and the software development. AA wrote the first draft of the manuscript, created the computational experiments and datasets, and created the first version of the `mergen` package. JAJ authored subsequent versions of `mergen` and added enhancements to the package. JAJ plotted the final versions of the figures and finalized the computational experiments. AM created the `mergenstudio` package with contributions from NAK and JAJ. All authors edited and finalized the manuscript.

